# Overexpressing eukaryotic elongation factor 1 alpha (eEF1A) proteins to promote corticospinal axon repair after injury

**DOI:** 10.1101/2022.04.14.488293

**Authors:** Daniel Romaus-Sanjurjo, Junmi M. Saikia, Hugo J. Kim, Kristen M. Tsai, Geneva Q. Le, Binhai Zheng

## Abstract

Although protein synthesis is hypothesized to have a pivotal role in axonal repair after central nervous system (CNS) injury, the role of core components of the protein synthesis machinery has not been examined. Notably, some elongation factors possess non-canonical functions that may further impact axonal repair. Here, we examined whether overexpressing eukaryotic elongation factor 1 alpha (eEF1A) proteins enhances the collateral sprouting of corticospinal tract (CST) neurons after unilateral pyramidotomy, along with the underlying molecular mechanisms. Compared with axonal regeneration from injured neurons, axonal sprouting from uninjured neurons occurs spontaneously after injury and may represent a more accessible form of axonal repair for clinical translation. We found that overexpressing eEF1A1, eEF1A2 or both proteins in CST neurons increased the levels of pS6, an indicator for mTOR activity, in neuronal somas. In contrast, the levels of pSTAT3 and pAKT were not increased. Strikingly, overexpressing eEF1A2 alone, but neither eEF1A1 alone nor both factors simultaneously, increased protein synthesis and actin rearrangement in CST neurons. While eEF1A1 overexpression only slightly enhanced CST sprouting across the midline into the denervated side in the cervical spinal after pyramidotomy, eEF1A2 overexpression substantially enhanced this sprouting. Surprisingly, co-overexpression of both eEF1A1 and eEF1A2 led to a sprouting phenotype similar to wild-type controls, suggesting an antagonistic effect of overexpressing both proteins. These data provide the first evidence that overexpressing a core component of the translation machinery, eEF1A2, enhances CST sprouting, likely by a combination of increased protein synthesis, mTOR signaling and actin cytoskeleton rearrangement.

## Introduction

Spinal cord injury (SCI) often causes an irreversible loss of function which is due in part to the damage of the corticospinal tract (CST). Since the CST contributes significantly to the control of skilled motor movements in humans, unraveling the mechanisms of axonal repair for the CST will benefit therapeutic development for SCI. Compared with the regeneration of injured axons, compensatory sprouting of uninjured axons represents a more accessible form of axonal repair for clinical translation due to three factors: the existence of spared axonal pathways in the majority of SCI patients, a higher level of intrinsic axon growth ability of uninjured neurons, and a lower barrier to attain and enhance sprouting^1^.

mRNA translation is a key process in cellular metabolism that leads to protein synthesis. During translation elongation, eEF1A proteins carry out the critical step of recruiting aminoacyl-tRNAs to the A site of the ribosome^2,3^. There are two eEF1A proteins, eEF1A1 and eEF1A2, encoded by two different genes with different expression patterns: eEF1A1 is present in all adult tissues but muscle and heart, while eEF1A2 is highly expressed in muscle and heart tissues as well as neurons^4,5^. Following axonal damage, the capacity of neurons to synthesize new proteins is crucial for their regenerative ability^6^. Indeed, the mTOR pathway is a key regulator of CNS axon regeneration^7,8^, partially by activating protein synthesis via the eukaryotic initiation factor 4E (eIF4E)^9,10^. Recently, manipulating eEF1A2 was found to enhance protein translation *in vitro* in a study of dendritic spine remodeling^11^. Given that both eEF1A1 and eEF1A2 carry out a pivotal function during protein synthesis, they are interesting targets to increase the rate of protein synthesis underlying axon growth.

In addition to their role in protein synthesis, eEF1A proteins also have non-canonical roles. They can promote cell growth and proliferation by serving as upstream activators of PI3K/AKT/mTOR or PI3K/AKT/STAT3 pathways^12–14^. Moreover, eEF1A proteins possess actin-binding domains that enable a role in actin-bundling and cytoskeleton organization *in vivo*^15–19^. This is relevant following axonal damage since actin dynamics have a key role in the formation of a competent growth cone, which is vital for proper axon growth and regeneration^20–22^. Overall, the literature has implicated a role for eEF1A proteins in cellular processes deemed important for axonal repair.

These considerations prompted us to test the possibility that manipulating eEF1A proteins through virally-mediated overexpression could enhance compensatory CST sprouting after a unilateral pyramidotomy injury by boosting protein translation and cytoskeleton rearrangement as well as activating downstream neuron-intrinsic pathways. Our data provide the first *in vivo* demonstration to our knowledge that manipulating a core component of the translational machinery can enhance axonal repair in the mammalian CNS.

## Materials and methods

### Mice

All mouse husbandry and experimental procedures were approved by the Institutional Animal Care and Use Committee at the University of California San Diego. A mix of wild type C57BL/6 male and female mice (6 weeks old) in an approximately 1:1 ratio were used for all experiments. In some procedures, these mice were compared against PTEN conditional knockout (cKO) mice in the same genetic background.

### Viral production

pCMV6-eEF1A1 (#MG207381) and pCMV6-eEF1A2 (#MG207396) mouse ORF clones (GFP tagged) were obtained from OriGene (USA). They were subcloned into AAV2 (shortened as AAV) vectors at the Boston Children’s Hospital Viral Core, from where AAV-Cre and AAV-GFP were also acquired (Suppl. Fig. 1). Viral titers were confirmed using qPCR to be 1.6×10^12^ TU/ml (eEF1A1-overexpressing AAV vector; eEF1A1 OE), 1.2×10^12^ TU/ml (eEF1A2-overexpressing AAV vector; eEF1A2 OE), and 0.5×10^12^ TU/ml (AAV-Cre and AAV-GFP). The viral titer of eEF1A1 OE was diluted to the one of eEF1A2 OE (1.2×10^12^ TU/ml) in order to have the same viral titer in the individual treatments. Likewise, this viral titer (1.2×10^12^ TU/ml) was maintained during the combination treatment for each construct (eEF1A1 OE + eEF1A2 OE).

### Surgical procedures

Virus was delivered following general anesthesia by intraperitoneal injection of ketamine and xylazine dosed at 80-100 mg/kg and 10 mg/kg, respectively. Briefly, a total of 1.2 μL of each vector separately (eEF1A1 OE and eEF1A2 OE), or co-injections of both vectors (eEF1As OE), or AAV-GFP as control were injected at 3 different sites (0.4 μL per site) into the right sensorimotor cortex of 6 week old wild-type mice^23^. Injection coordinates relative to bregma were as follows: 1.2 mm lateral, 0.5 mm anterior, 1.2 mm lateral, 0.5 mm posterior, 2.2 mm lateral, 0.0 mm anterior. For each injection site, the needle was lowered to a depth of 0.7 mm. PTEN^fl/fl^ mice underwent the same procedures with AAV-Cre to generate PTEN cKO mice, or with AAV-GFP as controls.

Unilateral pyramidotomy and biotinylated dextran amine (BDA) tracer injection were performed as described previously^23–25^. Mice were sacrificed at 12 weeks of age (Suppl. Fig. 1).

### In vivo puromycin administration and tissue processing

A group of mice were injected intraperitoneally with 225 mg/kg puromycin (Thomas Scientific, C791P84) as previously described^26^. For all experiments, mice were sacrificed by an overdose of Fatal Plus (pentobarbital sodium, intraperitoneal injection of 150mg/kg mouse), and then transcardially perfused with an ice-cold solution of 4% paraformaldehyde (PFA). Finally, brains and spinal cords were dissected out, and the tissues were post-fixed overnight at 4°C in the same fixative solution.

Tissue processing was performed as described previously^23,24^. Briefly, tissues were transversely sectioned with a cryostat at a thickness of 20 μm. For tissues containing BDA labeled axons, cervical spinal cord and medulla sections were incubated in Vectastain ABC solution (Vector Laboratories) overnight at 4°C. BDA was detected with TSA Plus Fluorescein System (1:200, PerkinElmer). For immunohistochemistry, incubation in the primary antibody solution (see concentrations below) was carried out at 4°C overnight. Sections were washed, followed by secondary antibody staining for 2 hr at room temperature (antibody solutions at 1:500). Next, sections were washed again, incubated with DAPI for 10 min, and mounted with Fluoromount-G (Southern Biotech).

The following antibodies were used: rabbit anti-eEF1A1 (1:100, abcam), rabbit anti-eEF1A2 (1:100, Sigma-Aldrich), monoclonal mouse anti-Puromycin, clone 12D10 (1:100, EMD Millipore), guinea pig anti-NeuN (1:200, Sigma-Aldrich), rabbit anti-pS6 (Ser235/236) (1:200, Cell Signaling Technology), rabbit anti-pSTAT3 (Tyr705) (1:200, Cell Signaling Technology), rabbit anti-pAKT1 (Ser473) (1:100, Cell Signaling Technology), rat anti-GFAP (1:500, Thermo Fisher), and rabbit anti-beta Actin (1:100, abcam). For sprouting studies, selected transverse sections of cervical spinal cord (C7) were immunostained for PKCγ (1:100, Santa Cruz Biotechnology) to examine the completeness of the lesion for each mouse^25^. Mice with incomplete lesion were excluded from the study.

For pSTAT3 and beta Actin staining that required antigen retrieval, the protocol was performed as described previously^27^. Then, sections were incubated with anti-pSTAT3 or anti-beta actin antibodies for overnight at RT, and then following the regular protocol.

For puromycin staining, the Mouse on Mouse (MOM) Detection Kit (Vector Labs, BMK-2202) was used for blocking and staining procedures as previously described^26^, with buffers prepared as described in standard protocol supplied with the kit.

### Image acquisition

Stained tissue sections were photographed using an upright epifluorescence microscope (Zeiss Axio Imager M1). For coronal brain sections, one section per injection site (3 sections total) showing GFP signal along layer V of the cerebral cortex were photographed for each animal. Five transverse sections of the medullary pyramids were randomly photographed. For cervical spinal cord sections, 10 transverse sections per animal between C5-C7 levels were randomly photographed. Photomicrographs were taken at 10× (brain, medulla and cervical sections) and 20× (brain sections) magnification without changing the amplifier gain or the offset to avoid the introduction of experimental variability. Following quantifications, contrast and brightness were minimally adjusted in figures, uniformly across panels for each experiment, with Adobe Photoshop CS4 (Adobe Systems).

### Immunofluorescence quantifications

To quantify the intensity of each immunofluorescence (IF) signal in layer V CST neurons, mean fluorescent intensity (mean grey value) from GFP+ CST neurons was determined using ImageJ. All values were normalized against control values. The experimenter was blinded during quantifications.

For the quantification of BDA-labeled axon numbers, we adapted a semiautomatic quantification method previously used^28^. Positive profiles of the medulla were quantified by using the “analyze particle” function of ImageJ, and the average number of BDA positive axons per section was calculated for each animal: 802.2 ± 35.83 (control); 739.9 ± 44.16 (eEF1A1 OE mice); 748.1 ± 43.35 (eEF1A2 OE mice); 825.3 ± 80.97 (eEF1As OE mice); and 652.9 ± 72.12 (PTEN cKO mice) (Fig. 7J; Suppl. Fig. 2E). Both the initial observation of the stained sections and later quantification of axon numbers were conducted by experimenters blinded to genotypes.

Sprouting Axon Number Index was quantified similarly as described^23,24^. Briefly, the number of axons crossing pre-defined lines at various distances from the midline on the denervated gray matter was manually counted in 10 randomly chosen sections per animal by a blinded observer. Counts were averaged for each animal and normalized against the total labeled CST axon count in the medulla^25^ to obtain the sprouting index, which was plotted as a function of distance from the midline.

### Statistical analyses

Statistical analysis was carried out using Prism 6 (GraphPad software, La Jolla, CA). Data were presented as mean ± S.E.M. The immunohistochemistry data with multiple comparisons were analyzed by either one-way ANOVA or Kruskal–Wallis test where appropriate, and post-hoc Dunn’s multiple comparisons test. The results of control vs. eEF1A overexpression groups were analyzed by Mann–Whitney U test. Sprouting index data were analyzed via two-way repeated measures ANOVA with Tukey post-hoc test. Correlations were assessed with the Spearman correlation coefficient test or the Pearson correlation coefficient test where appropriate. The significance level was set at 0.05. In the figures, significance values were represented by different number of asterisks: \**p*<0.05; \*\**p*<0.01; \*\*\**p*<0.001; \*\*\*\**p*<0.0001. Specific n values (cells and/or animals) for each study are listed in figure legends.

## Results

### Cortical injections of AAV-eEF1A1 or AAV-eEF1A2 increase the expression of eEF1A1 and eEF1A2 proteins respectively in CST neurons

Prior to testing our hypothesis, we first validated the effect of injecting AAV-eEF1A1 or AAV-eEF1A2 in CST neurons of layer V cortex. These CST neurons are identifiable based on their larger size compared to other counterparts, positive staining for NeuN and GFP, and anatomical location in cortical layer V. In eEF1A1 OE mice, eEF1A1 immunoreactivity in GFP^+^ CST neurons was increased by 29.2% as compared with control mice (Mann-Whitney *U, p*<0.0001; Fig. 1A-B’’’’, E). Following eEF1A2 overexpression, GFP^+^ CST neurons showed a significant 38.5% increase in the eEF1A2 immunoreactivity compared with controls (Mann-Whitney *U, p*<0.0001; Fig. 1C-D’’’’, F). Consistent with the GFP-fusion expression constructs, the GFP signals from either AAV-eEF1A1::GFP or AAV-eEF1A2::GFP co-labeled with elevated eEF1A1 and eEF1A2 signals, respectively (Fig. 1A’’’, C’’’; Suppl. Fig. 1). These data confirmed that each viral vector led to the overexpression of eEF1A1 or eEF1A2 respectively in CST neurons.

**Figure 1.**
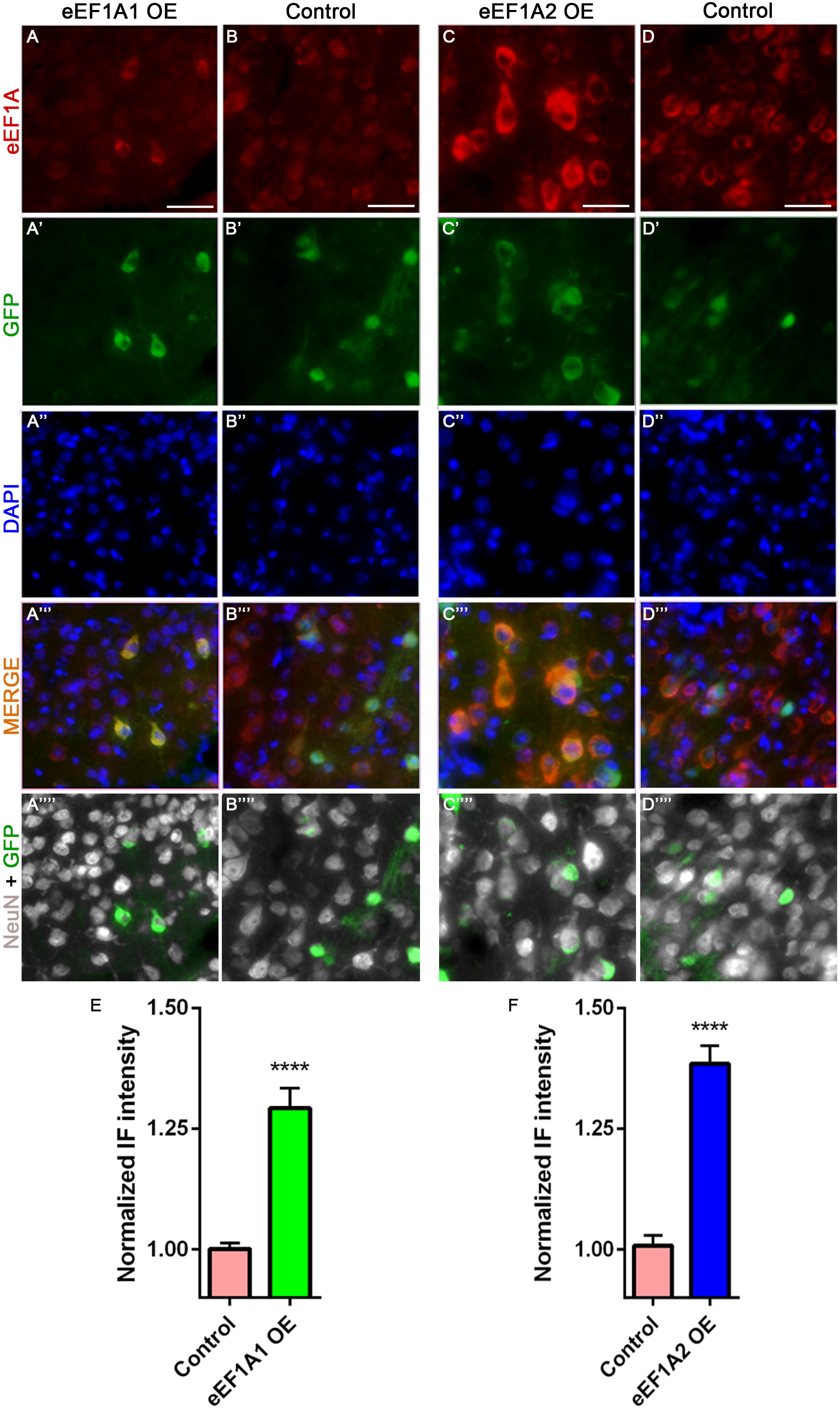
Infection by AAV-eEF1A1 or AAV-eEF1A2 increased the level of eEF1A1 and eEF1A2 proteins, respectively, in CST neurons. Representative images of eEF1A1, GFP (viruses), and NeuN staining at the layer V of the right sensorimotor cortex of injured eEF1A1-overexpressing (OE) mice (A-A’’’’); and injured control mice (AAV-GFP injection) (B-B’’’’). Representative images of eEF1A2, GFP (viruses), and NeuN staining at the layer V of the right sensorimotor cortex of injured eEF1A2-OE mice (C-C’’’’); and injured control mice (AAV-GFP injection) (D-D’’’’). E-F: Quantification of eEF1A1 and eEF1A2 immunoreactivity. Scale bars: 50 μm. Control, 12 mice; eEF1A1, 10 mice; eEF1A2, 10 mice; 20 cells quantified per mouse. Stats: D’Agostino Normality Test; Mann–Whitney U test. Bars show mean ± SEM. ****p<0.0001.

### Overexpressing eEF1A2 but not eEF1A1 increases protein synthesis in CST neurons

Given that eEF1A proteins play a pivotal role during mRNA translation^2,11^, we examined the effect of overexpressing either eEF1A1 or eEF1A2 on protein synthesis in CST neurons (Fig. 2). We used a puromycin-based assay, Surface Sensing of Translation (SUnSET), previously adapted for use *in vivo*^26,29^. The SUnSET assay showed that eEF1A1 overexpression did not significantly increase protein synthesis in CST neurons (Fig. 2B-B’’’’, F). In contrast, overexpressing eEF1A2 significantly albeit modestly increased protein synthesis (by ∼5.6%) compared to control (Kruskal–Wallis, *p*=0.0398; Fig. 2C-C’’’’, F). Unexpectedly, co-overexpressing both eEF1A1 and eEF1A2 simultaneously led to a slight trend for increased protein synthesis that did not reach statistical significance (Fig. 2D-D’’’’, F). We used PTEN cKO mice as a positive control since PTEN deletion is known to enhance CST regeneration and the mTOR pathway regulates global translation^9^. Unsurprisingly, these mice exhibited a more substantial ∼22.3% increase in protein translation (Kruskal–Wallis, *p*<0.0001; Fig. 2E-E’’, F). Together, these results revealed that only the overexpression of eEF1A2 led to a significant increase in mRNA translation, which was blocked by co-overexpressing eEF1A1.

**Figure 2.**
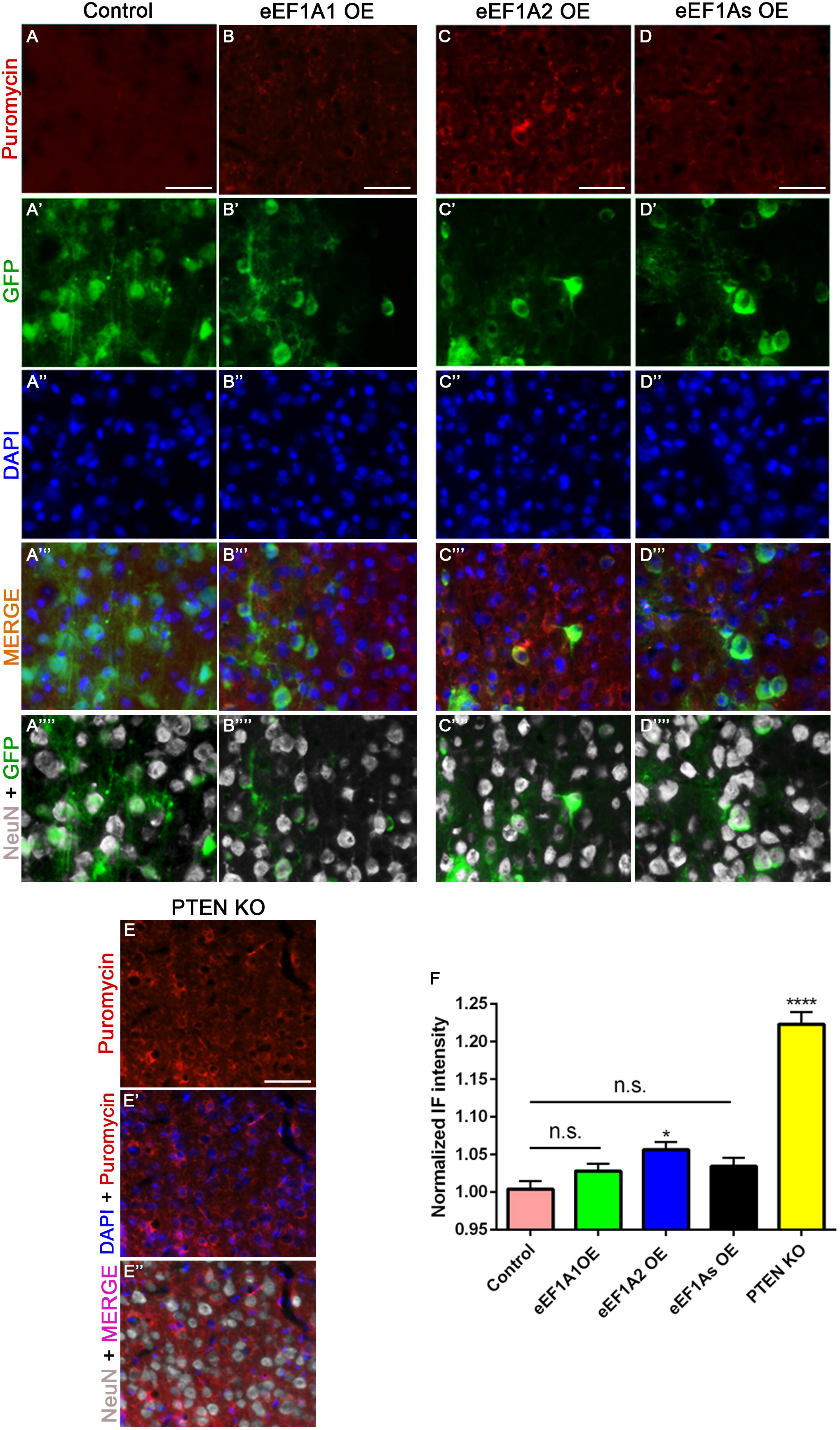
Only eEF1A2 OE mice increase protein synthesis. Representative images of puromycin, GFP (viruses), and NeuN staining at the layer V of the right sensorimotor cortex of injured control mice (AAV-GFP injection) (A-A’’’’); injured eEF1A1-overexpressing (OE) mice (B-B’’’’); injured eEF1A2-OE mice (C-C’’’’); injured eEF1As-OE mice (D-D’’’’); and injured PTEN cKO mice serving as a positive control (E-E’’). F: Quantification of puromycin immunoreactivity. Scale bars: 50 μm. 2 mice per each genetic condition; 60 cells quantified per mouse. Stats: D’Agostino Normality Test; one-way ANOVA with Kruskal-Wallis Test. Bars show mean ± SEM. *p=0.0398, ****p<0.0001.

### Overexpressing eEF1A1 and eEF1A2 impacts mTOR and STAT3 pathways differently in CST neurons

eEF1A proteins can act as upstream activators of mTOR and STAT3 pathways in non-neuronal cells^12,13^, both of which are well known regulators of CST axon sprouting after CNS injury^7,30^. Immunohistochemistry revealed an increase in the levels of pS6, a marker for mTOR activity, following eEF1A1 OE (33.9%), eEF1A2 OE (37%) or co-overexpression (33.1%) in CST neurons compared to controls (Kruskal–Wallis, *p*<0.0001 for all; Fig. 3). As a positive control, PTEN deletion increased pS6 levels by a large 106% as compared to controls (Kruskal–Wallis, *p*<0.0001; Fig. 3E-E’’, F). On the other hand, no significant differences were found in pSTAT3 immunostaining among mice overexpressing eEF1A1, eEF1A2 both proteins and control mice (Kruskal–Wallis; Fig. 4). Together, these results showed that overexpressing eEF1A proteins alone or in combination moderately activates the mTOR pathway as previously seen in non-neuronal cells, but had no detectable effect on the STAT3 pathway.

**Figure 3.**
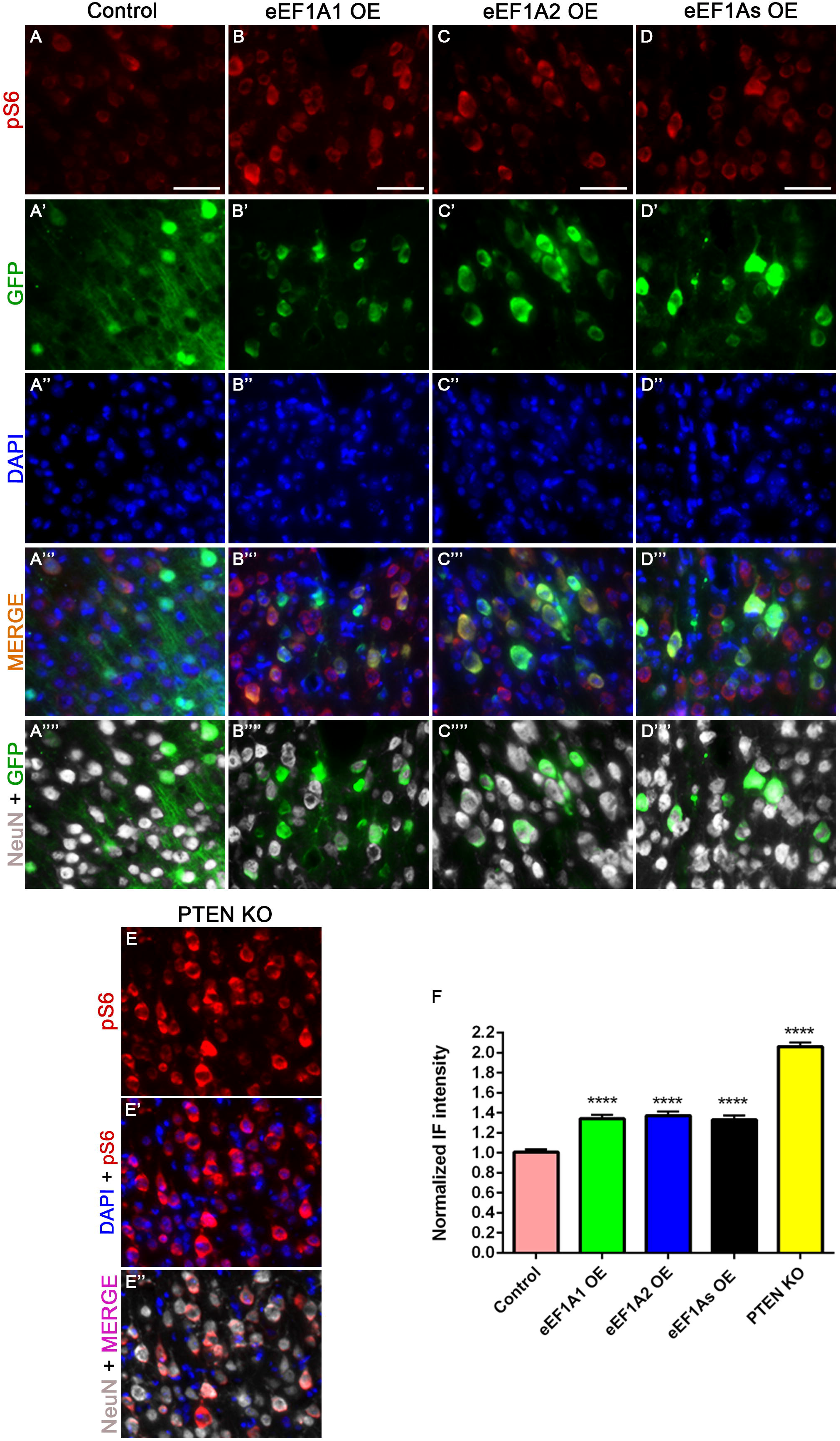
Overexpression of eEF1A proteins activates mTOR signaling. Representative images of pS6, GFP (viruses), and NeuN staining at the layer V of the right sensorimotor cortex of injured control mice (AAV-GFP injection) (A-A’’’’); injured eEF1A1-overexpressing (OE) mice (B-B’’’’); injured eEF1A2-OE mice (C-C’’’’); injured eEF1As-OE mice (D-D’’’’); and injured PTEN cKO mice serving as a positive control (E-E’’). F: Quantification of pS6 immunoreactivity. Scale bars: 50 μm. Control, 12 mice; eEF1A1, 10 mice; eEF1A2, 10 mice; eEF1As, 8 mice; PTEN cKO, 4 mice; 20 cells quantified per mouse. Stats: D’Agostino Normality Test; one-way ANOVA with Kruskal-Wallis Test. Bars show mean ± SEM. ****p<0.0001.

**Figure 4.**
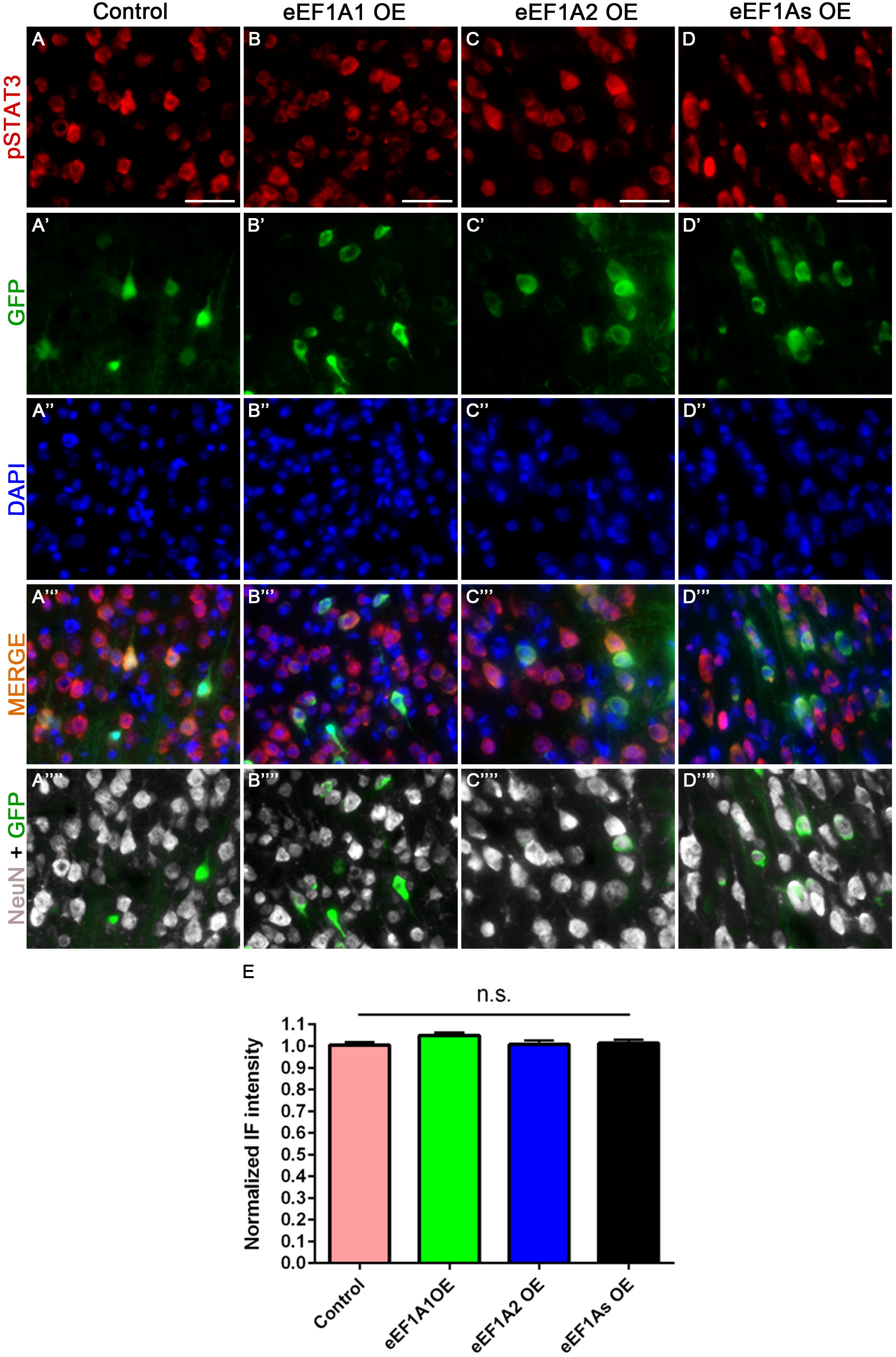
Overexpression of eEF1A proteins does not activates STAT3 signaling. Representative images of pSTAT3, GFP (viruses), and NeuN staining at the layer V of the right sensorimotor cortex of injured control mice (AAV-GFP injection) (A-A’’’’); injured eEF1A1-OE mice (B-B’’’’); injured eEF1A2-OE mice (C-C’’’’); and injured eEF1As-OE mice (D-D’’’’). E: Quantification of pSTAT3 immunoreactivity. Scale bars: 50 μm. Control, 12 mice; eEF1A1, 10 mice; eEF1A2, 10 mice; eEF1As, 8 mice; PTEN cKO, 4 mice per genetic condition; 20 cells quantified per mouse. Stats: D’Agostino Normality Test; one-way ANOVA with Kruskal-Wallis Test revealed no significant differences. Bars show mean ± SEM.

### Overexpressing eEF1A2 but not eEF1A1 promotes actin rearrangement

eEF1A proteins possess actin-binding domains that enable a role in actin-bundling and cytoskeleton organization *in vivo* and *in vitro*^15,16^. Thus, we examined the impact of overexpressing eEF1A proteins on actin remodeling. Whereas control, eEF1A1 OE and eEF1As OE mice displayed a diffuse pattern of beta Actin expression (Fig. 5A); this pattern appeared bundled in eEF1A2 OE mice, defining the cellular shape (Fig. 5B, arrowheads). This qualitative observation was corroborated with immunohistochemical quantifications, which revealed a significant increase of actin signal intensity only in eEF1A2 OE mice (one-way ANOVA, *p* = 0.0004; Fig. 5C). Interestingly, a few GFP^-^ /GFAP^+^ astrocytes (Fig. 5, thick arrows) and many GFP^-^/GFAP^-^/NeuN^-^ cells (Fig. 5, thin arrows) from eEF1A2 OE mice also showed bundled beta actin expression. Overall, this raises the possibility that the overexpression of eEF1A2 alters actin dynamics by bundling actin not only directly in neurons but also indirectly in non-neuronal cells through an unknown neuron-glia crosstalk mechanism.

**Figure 5.**
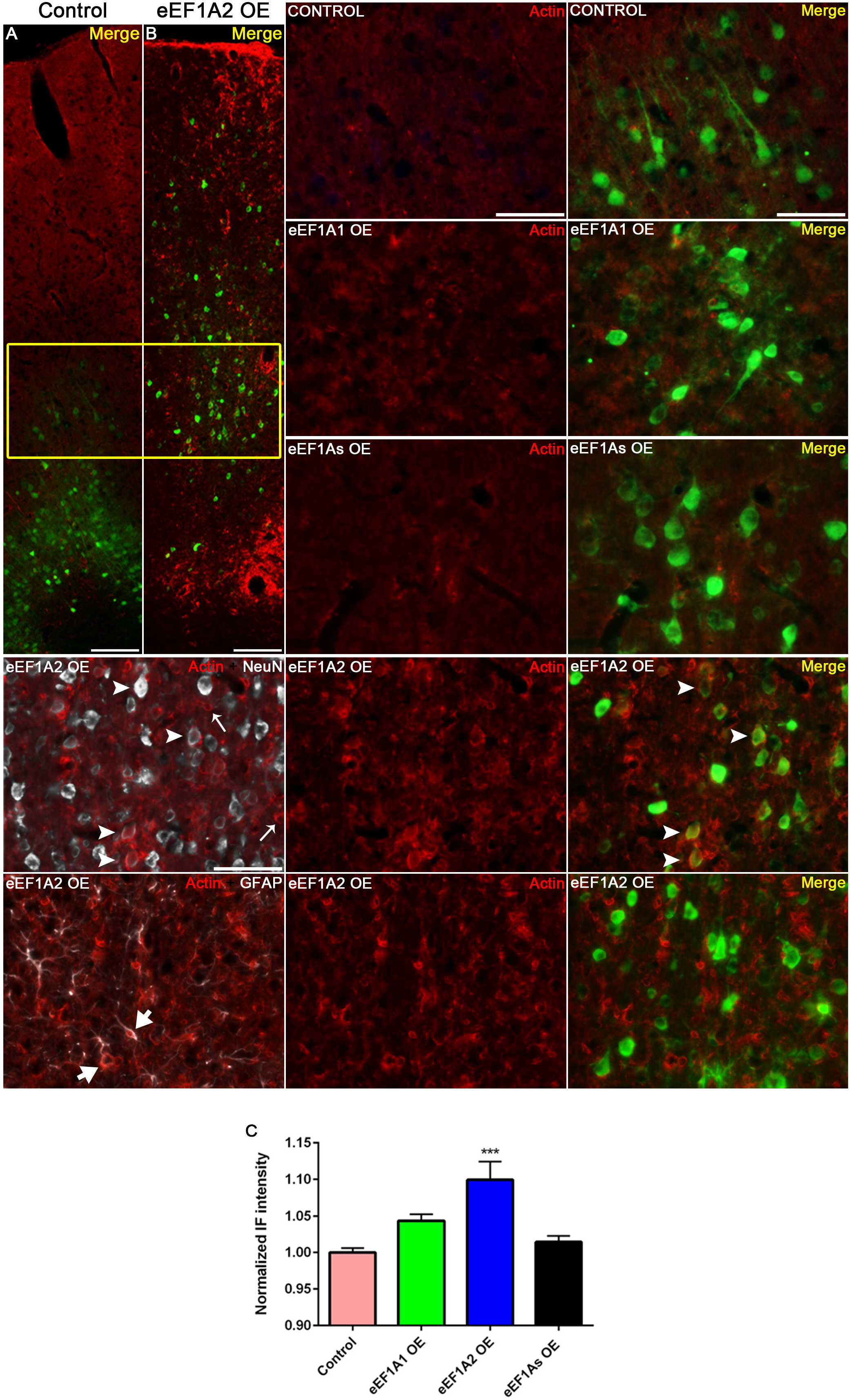
Only eEF1A2 OE mice display actin rearrangement. Left panels: Representative 10× images of β-actin and GFP (viruses) staining at the right sensorimotor cortex of in control (A) and eEF1A2 OE (B) mice. Right and bottom panels: Detail from the cortical layer V (yellow box) at 20× magnification. Please note the bundled pattern of β-actin staining defining the cellular shape in NeuN^+^ cells (arrowheads), GFP^-^/GFAP^+^ cells (thick arrows), and GFP^-^/GFAP^-^/NeuN^-^ cells (thin arrows). Scale bars = 50 µm. C: Quantification of β-actin immunoreactivity. 3 mice per genetic condition. Stats: Kolmogorov-Smirnov Normality test; ordinary one-way ANOVA. Bars show mean ± SEM. ***p<0.0004.

### Neither eEF1A1 overexpression nor eEF1A2 overexpression increased the level of pAKT in CST neurons

Given that AKT is involved in both the activation of the mTOR signaling^31^ and actin remodeling^15^, we examined the levels of AKT activation in our mTOR and beta actin results. The overexpression of eEF1A proteins did not significantly increase the levels of AKT phosphorylation (Kruskal Wallis; Fig. 6). Only PTEN KO mice revealed a statistically significant albeit modest 4% increase of pAKT signals (Kruskal–Wallis, *p* = 0.0402; Fig. 6E-E’’, F). Therefore, we cannot resolve whether eEF1A-mediated mTOR activation or eEF1A2-mediated actin rearrangement goes through AKT signaling.

**Figure 6.**
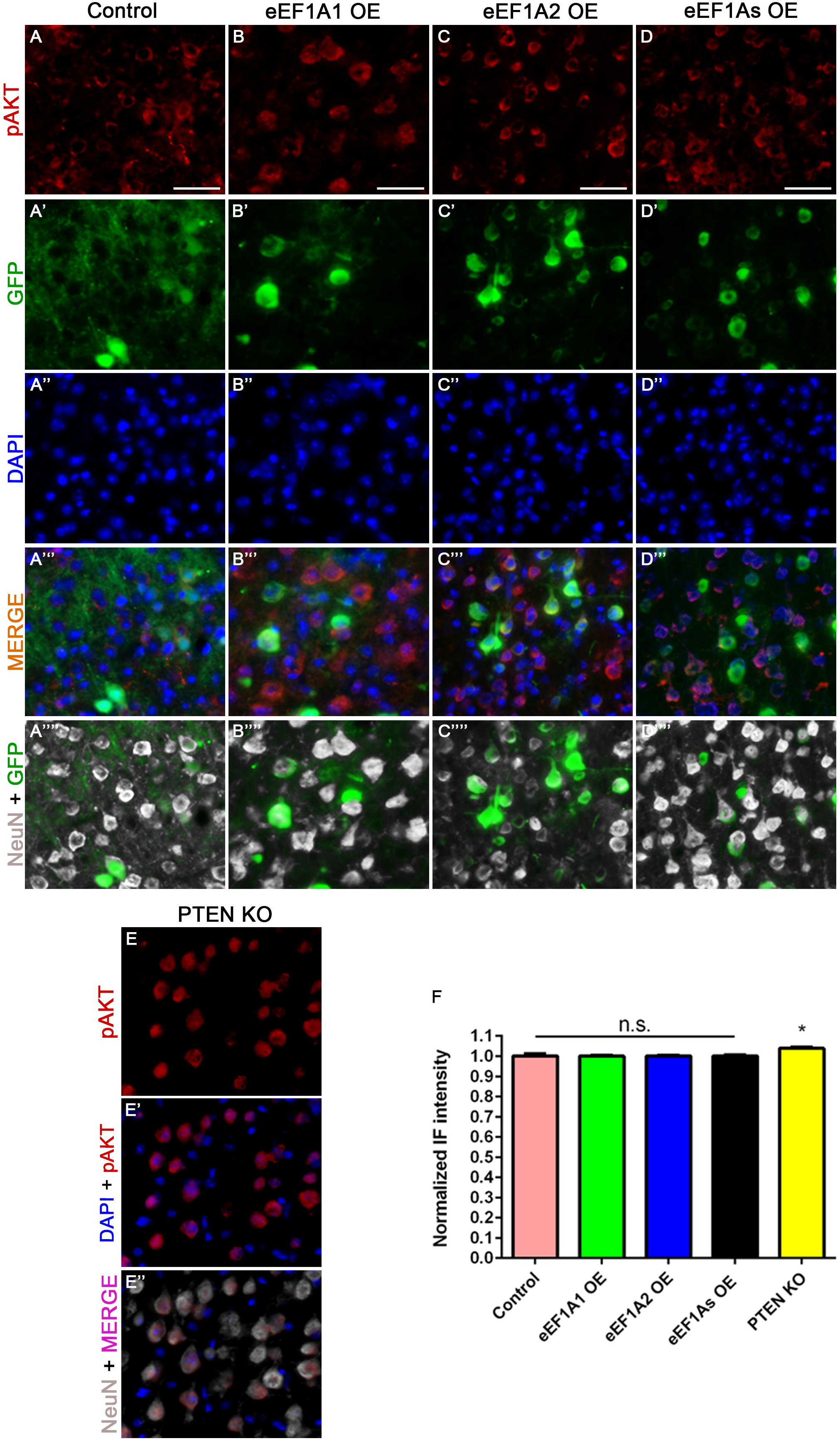
Overexpression of eEF1A proteins does not increase pAKT phosphorylation. Representative images of pAKT, GFP (viruses), and NeuN staining at the layer V of the right sensorimotor cortex of injured control mice (AAV-GFP injection) (A-A’’’’); injured eEF1A1-overexpressing (OE) mice (B-B’’’’); injured eEF1A2-OE mice (C-C’’’’); injured eEF1As-OE mice (D-D’’’’); and injured PTEN cKO mice (E-E’’). F: Quantification of pAKT immunoreactivity. Scale bars: 50 μm. 3 mice per each genetic condition; 50 cells quantified per mouse. Stats: D’Agostino Normality Test; one-way ANOVA with Kruskal-Wallis Test. Bars show mean ± SEM. *p=0.0402.

### Overexpressing eEF1A2 promotes CST sprouting more than eEF1A1

After undergoing unilateral pyramidotomy (Fig. 7A-D, I-J), eEF1A1 OE mice only exhibited a significant increase in the number of axons just crossing the midline at 50 µm compared to GFP control mice (two-way repeated measures [RM] ANOVA with Tukey post hoc test, p = 0.0106) (Fig. 7B, I). In contrast, eEF1A2 OE mice exhibited significantly elevated levels of CST sprouting up to 200 µm past the midline (two-way RM ANOVA with Tukey’s correction; p = 0.0006 at 50 µm; p = 0.0101 at 100 µm; and p = 0.0044 at 200 µm) (Fig. 7C, I). Moreover, the level CST sprouting in eEF1A2 OE mice was significantly higher than the one in eEF1As mice at 200 µm (two-way RM ANOVA with Tukey’s correction; p = 0.0403; Fig. 7I). Interestingly, the sprouting indices from eEF1A2 OE mice were similar to those from PTEN cKO mice, where the two groups did not exhibit statistically significant differences (two-way RM ANOVA with Tukey’s correction) (Suppl. Fig. 2). The co-overexpression of eEF1A proteins unexpectedly suppressed this enhancement in CST sprouting (Fig. 7A, D, I).

**Figure 7.**
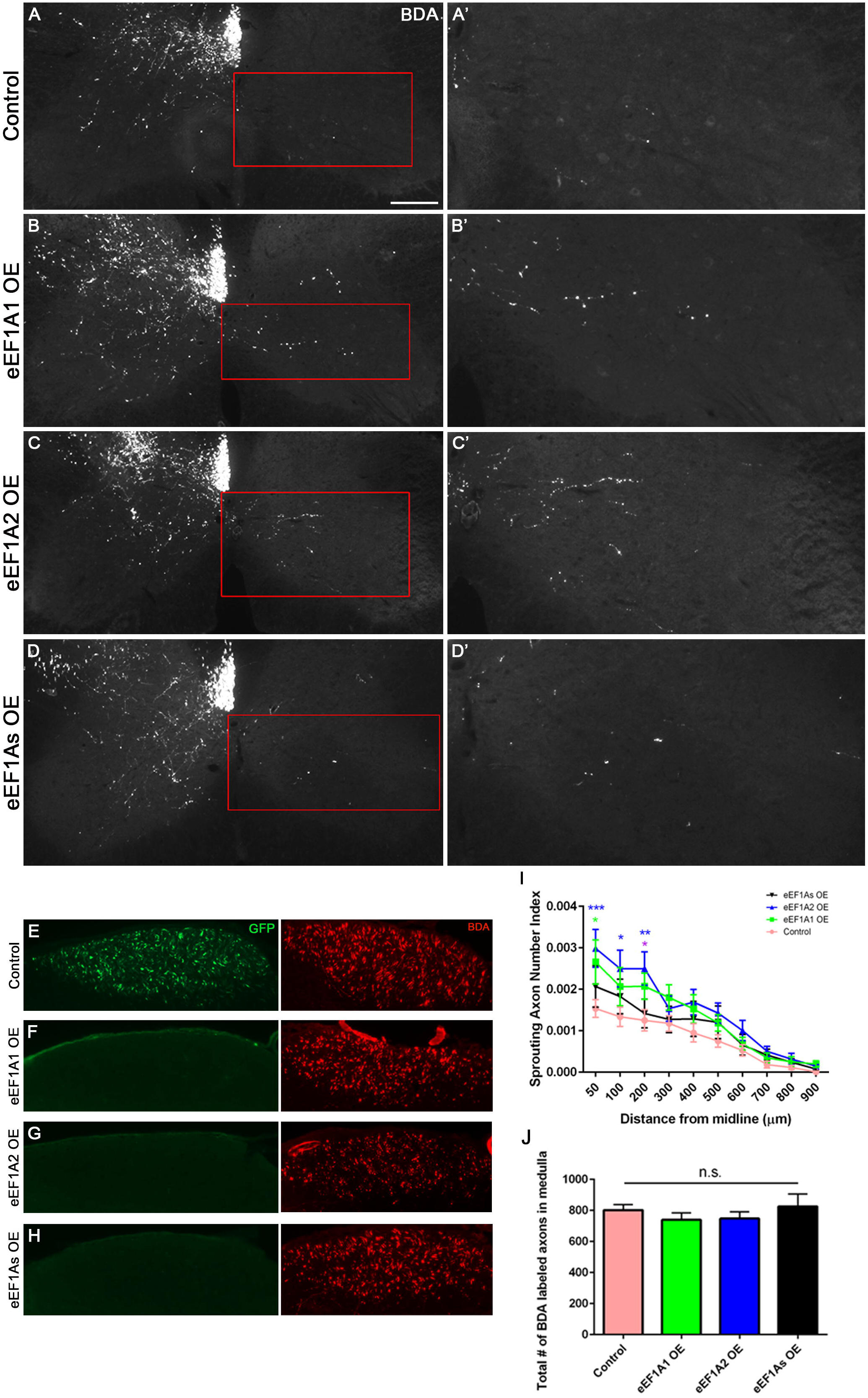
eEF1A2 OE exhibits increased compensatory sprouting more than eEF1A1 OE. Representative images of BDA tracing at the level of cervical spinal cord in control (A-A’), eEF1A1 OE (B-B’), eEF1A2 OE (C-C’), and eEF1As OE (D-D’) mice; scale bar = 300 µm. Right panel shows individual sprouting axons in the grey matter of the denervated half of spinal cord. E-H: Representative images of GFP and BDA axonal labeling at the level of medullas. I: Quantification of eEF1A1 OE, eEF1A2 OE, and eEF1As OE sprouting. Sprouting index indicates the ratio of the average number of axons counted at each distance past midline relative to the number of fibers labeled in the medulla. Compared to control, two-way RM ANOVA multiple comparisons with Tukey’s correction revealed elevated sprouting for eEF1A1 OE: at 50 µm, p = 0.0059. Elevated sprouting for eEF1A2 OE compared to controls was observed at 50 µm, p = 0.0003; at 100 µm, p = 0.0056; and at 200 µm, p = 0.0024; compared to eEF1A1 OE mice, sprouting in eEF1A2 OE was higher at 200 µm, p = 0.0487. J: Quantification of BDA-labeled axons at medullas. One-way ANOVA revealed no significant differences. Bars show mean ± SEM. Control mice: n = 12; eEF1A1 OE mice: n = 10; eEF1A2 OE mice: n = 9; eEF1As OE mice: n = 8; and PTEN cKO mice: n = 4.

Intriguingly, control mice with AAV-GFP injections exhibited GFP signals in the medulla, where mice injected with the AAV-eEF1A1::GFP or AAV-eEF1A2::GFP fusion construct had no detectable GFP signals (Fig. 7E-H, Suppl. Fig. 1). As a comparison, all mice exhibited robust BDA signals at the medulla level (Fig. 7E-H). This result suggests that the fusion proteins did not reach the axons at detectable levels. Thus, any overexpressed eEF1A protein likely acted in the somas and not axons (or dendrites) to enhance CST axon sprouting as might have been expected from the literature^11,32,33^. Together, these data indicate that overexpressing eEF1A1 and even more so eEF1A2 enhances CST sprouting through a mechanism in the neuronal somas.

## Discussion

In this study, we have provided gain of function data showing that virally-mediated overexpression of eEF1A proteins and especially eEF1A2 enhances axon sprouting of CST neurons following a unilateral pyramidotomy injury. This represents the first proof-of-concept demonstration that forced expression of a core component of the protein translation machinery can exert a beneficial effect on axonal repair after CNS injury *in vivo*. In the case of eEF1A2, this could occur by a combination of enhancing protein synthesis, activating the mTOR pathway, and promoting actin remodeling. Here we discuss the implications and possible mechanisms underlying this eEF1A-mediated enhancement of CST axon sprouting.

The *in vivo* puromycin incorporation assay showed that overexpressing eEF1A2 but not eEF1A1 significantly increased protein synthesis in CST neurons. While this increase was modest at ∼5.6%, such a modest increase in global protein synthesis can have a widespread effect on cellular metabolisms^11^. Importantly, recent work from Mendoza and colleagues^11^ showed that eEF1A2 can stimulate protein translation *in vitro*, which agrees with our *in vivo* results in the neuronal soma. The lack of detectable effect from eEF1A1 overexpression on translation may initially be surprising, but it may reflect the well-documented but incompletely understood developmental switch from eEF1A1 to eEF1A2 in neurons and their different biochemical properties^4,34–37^. Indeed, the lack of eEF1A2 in the postnatal stage leads to early neurodegeneration, muscle wasting, and subsequent animal death^38^. Overall, it aligns with our results where the extra eEF1A1 protein has no effect on neuronal protein synthesis, and even attenuates the effect of eEF1A2 on protein synthesis when both proteins are co-overexpressed in adult mice. Biochemically, eEF1A1 and eEF1A2 differ in their affinity for GDP/GTP^39,40^ but also in their ability to interact with Ca^2+^-calmodulin, which binds to eEF1A1 and displaces the tRNA^36^. Neurons are exposed to rapid changes in [Ca^2+^] and, so, this may promote the interaction eEF1A1-calmodulin and impede protein synthesis.

From cancer studies, it was established that eEF1A proteins can promote cell growth and proliferation by serving as an upstream activator of PI3K/AKT/mTOR^12,41^ and/or PI3K/AKT/STAT3^13,14^ pathways. In partial agreement with those *in vitro* studies, our results showed that in an *in vivo* model of CNS injury, the overexpression of eEF1A proteins elevated mTOR activity, whereas co-overexpressing both proteins had no synergistic effect. However, the overexpression of neither eEF1A protein promoted activation of the STAT3 pathway. Several *in vitro* studies have shown the ability of eEF1A proteins to either enhance or attenuate the activity of various kinases such as AKT and PKR by direct interaction^15,40,42–44^, although the underlying mechanism is still to be elucidated. Thus, it is possible that the overexpression of eEF1A proteins in CST neurons enhances the interactions between these elongation factors and such kinases along the mTOR pathway. Future studies will be needed to decipher the exact mechanism of action for eEF1A proteins in neuronal signaling.

The non-canonical role of eEF1A proteins related to actin dynamics has been most studied in the literature, mainly in yeast^15,16,19,45,46^. Importantly, *in vitro* studies confirmed that mammalian eEF1A proteins maintain the ability to interact with actin^11,36,47^. Here, we observed apparent bundling of beta actin *in vivo* in eEF1A2 OE mice, in contrast to control and eEF1A1 OE mice. These differences can be explained by a previous *in vitro* study reporting that mammalian eEF1A proteins differ in their capacity to interact with and modulate actin cytoskeleton^36^. Moreover, our results showed that additionally co-overexpressing eEF1A1 attenuates the actin rearrangement seen in eEF1A2 OE mice. It is known that eEF1A1 and eEF1A2 can interact with one another and co-localize in the cytoplasm^48^, so this result suggests that eEF1A1 interacts with eEF1A2, thereby interfering with its interaction with actin.

AKT kinase is a common upstream signaling molecule shared by actin rearrangement and mTOR pathways^15,49–51^. We speculated that the eEF1A2-mediated elevation of both mTOR activity and actin bundling would be associated with a higher level of phosphorylated AKT. However, we did not detect elevated levels of pAKT following eEF1A1 or eEF1A2 OE. Recently, Huang and colleagues^50^ reported that despite the important role of AKT in PTEN deletion-induced retinal axon regeneration, AKT activation is only marginal even in PTEN cKO mice, due to a mTORC1/S6K1-mediated feedback inhibition. This agrees with our results and can explain that only PTEN cKO mice exhibited a slight but significant increase in pAKT. Based on the pAKT results, we cannot conclude whether the activity of actin bundling seen in eEF1A2 OE mice is primarily mediated by direct interaction between eEF1A2 and actin, or indirectly through AKT signaling. Further studies are needed to decipher the precise interaction between eEF1A proteins and the different kinases underlying cell growth and cytoskeleton dynamics.

Finally, whereas eEF1A1 OE mice showed a modest enhancement of CST sprouting, eEF1A2 OE led to a clearer enhancement of CST sprouting. The pattern of effects on protein synthesis, mTOR signaling and actin dynamics support the hypothesis that eEF1A2 promotes sprouting through a combination of these three aspects rather than mTOR signaling alone, where the effects of eEF1A1 and eEF1A2 are indistinguishable. Accordingly, results from eEF1A2 OE mice suggest a higher rate in actin bundling, which may facilitate the generation of collaterals to innervate the denervated side^52^, akin to what has been described on dendritic spines^11^. In this regard, our study echoes strongly with the study by Mendoza and colleagues^11^ in that eEF1A2 serves as a link to coordinate protein translation and actin dynamics in regulating neuronal functions. Whereas Mendoza studied this link in the context of structural plasticity in dendritic spines, our data implicate this link in the context of axonal regeneration after CNS injury. Intriguingly, the interaction between both eEF1A proteins in eEF1As OE mice blocked the increased sprouting seen in eEF1A2 OE mice, which was associated with a lack of increase in protein synthesis and actin rearrangement. Furthermore, our results unexpectedly showed that the enhancement in CST sprouting is likely driven from the neuronal somas rather than axons^6,52–54^ since neither eEF1A1::GFP nor eEF1A2::GFP fusion proteins are localized to the level of medulla in CST axons.

## Conclusion

Together, our findings indicate that neuronal eEF1A2 promotes axon sprouting after CNS injury, and suggest that it does so by acting as a hub connecting mTOR signaling, protein synthesis and actin cytoskeleton. The level of eEF1A2-mediated enhancement in CST sprouting approached but did not exceed PTEN deletion. Our study provides the first demonstration that manipulating a core component of the translational machinery can enhance axonal sprouting in the mammalian CNS, supporting the hypothesis that protein synthesis is important for axonal repair. Unexpectedly, eEF1A2 appears to carry out these functions from the neuronal somas and not at the axons, while eEF1A1 exerts a somewhat antagonistic role that remains to be fully understood. Future studies will be required to determine whether the overexpression of eEF1A2 can also promote axon regeneration from injured CST neurons following SCI.

## Acknowledgements

This work has been Supported by grants from NIH/NINDS (NS093055), VA (RX002483), Wings for Life and Craig H. Neilsen foundations to B.Z. Viral preps were obtained from Boston Children’s Hospital Viral Core (EY012196). The contents do not represent the views of the U.S. Department of Veterans Affairs or the United States Government.

## Conflict of interest

The authors declare that the research was conducted in the absence of any commercial or financial relationships that could be construed as a potential conflict of interest.

**Supplementary Figure 1**. A: Experimental timeline and injury model illustration. B: Different viral constructs used in this study. C: Positive correlation for eEF1A1 and eEF1A2 immunofluorescence (IF) signals, and GFP IF signals in eEF1A1 OE and eEF1A2 OE mice, respectively (eEF1A1 IF in control mice: r =0.128, p = 0.0981, and in eEF1A1 OE mice: r =0.5381, p < 0.0001; eEF1A2 IF in control mice: r = -0.1084, p = 0.2712, and in eEF1A2 OE mice: r =0.2993, p < 0.0001).

**Supplementary Figure 2. eEF1A2 OE exhibits compensatory sprouting similar to PTEN cKO mice**. Representative images of BDA tracing at the level of cervical spinal cord in control (A-A’), PTEN KO (B-B’), and eEF1A2 OE (C-C’) mice; scale bar = 300 µm. Right panel shows individual sprouting axons in the grey matter of the denervated half of spinal cord. D: Quantification of PTEN cKO and eEF1A2 OE sprouting index. Compared to control, two-way RM ANOVA multiple comparisons with Tukey’s correction revealed elevated sprouting for PTEN cKO: at 50 µm, p < 0.0001; at 100 µm, p = 0.0263; and at 300 µm, p = 0.0482; and at 400 µm, p = 0.007. Elevated sprouting for eEF1A2 OE compared to controls was observed at 50 µm, p = 0.0001; at 100 µm, p = 0.0031; and at 200 µm, p = 0.0013. E: Quantification of BDA-labeled axons at medullas. One-way ANOVA revealed no significant differences. Bars show mean ± SEM. Control mice: n = 12; eEF1A2 OE mice: n = 9; and PTEN cKO mice: n = 4.

