## Supplementary figures and images for "Overexpressing eukaryotic elongation factor 1 alpha (eEF1A) proteins to promote corticospinal axon repair after injury"

### Supplemental Figure 1

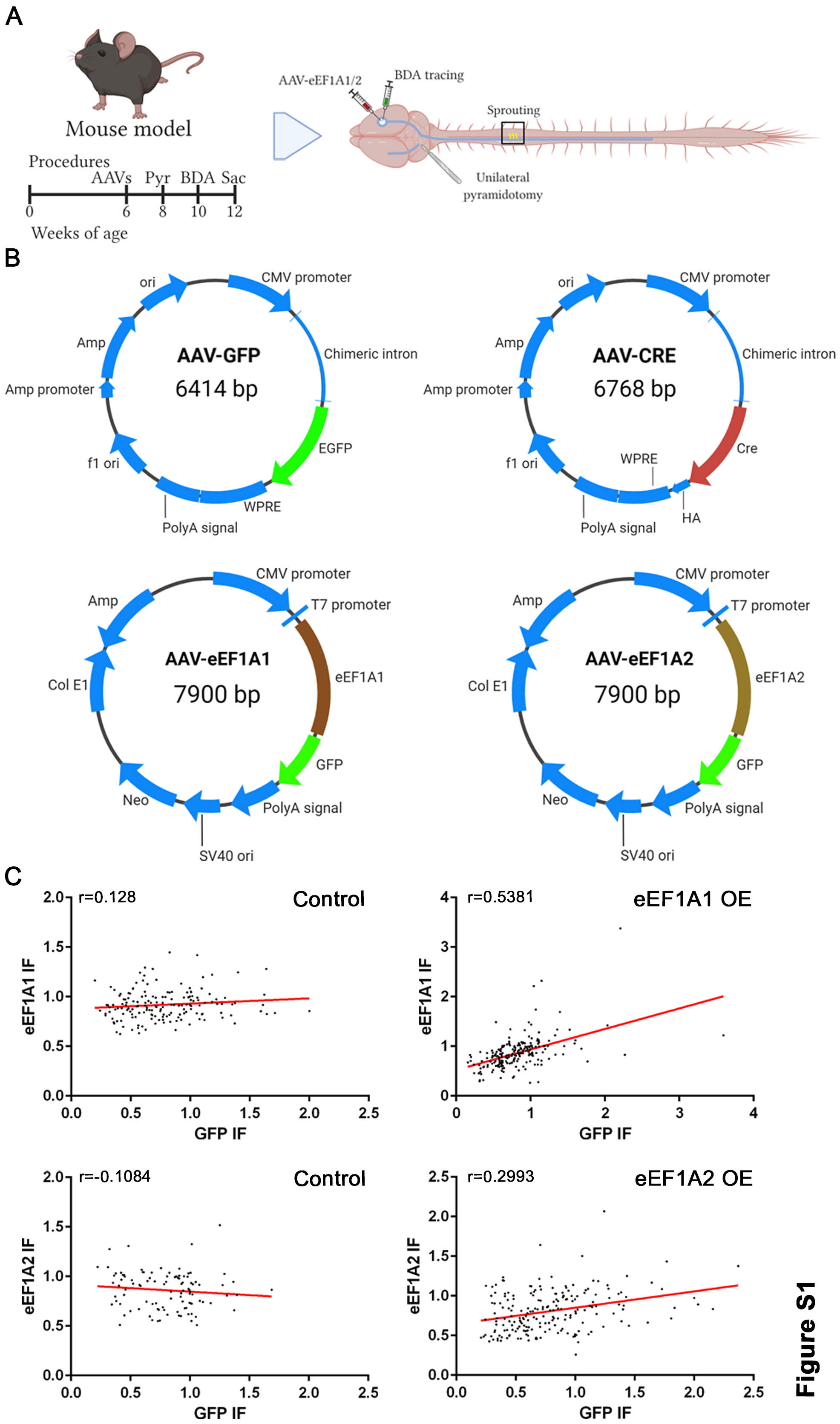

### Supplemental Figure 2

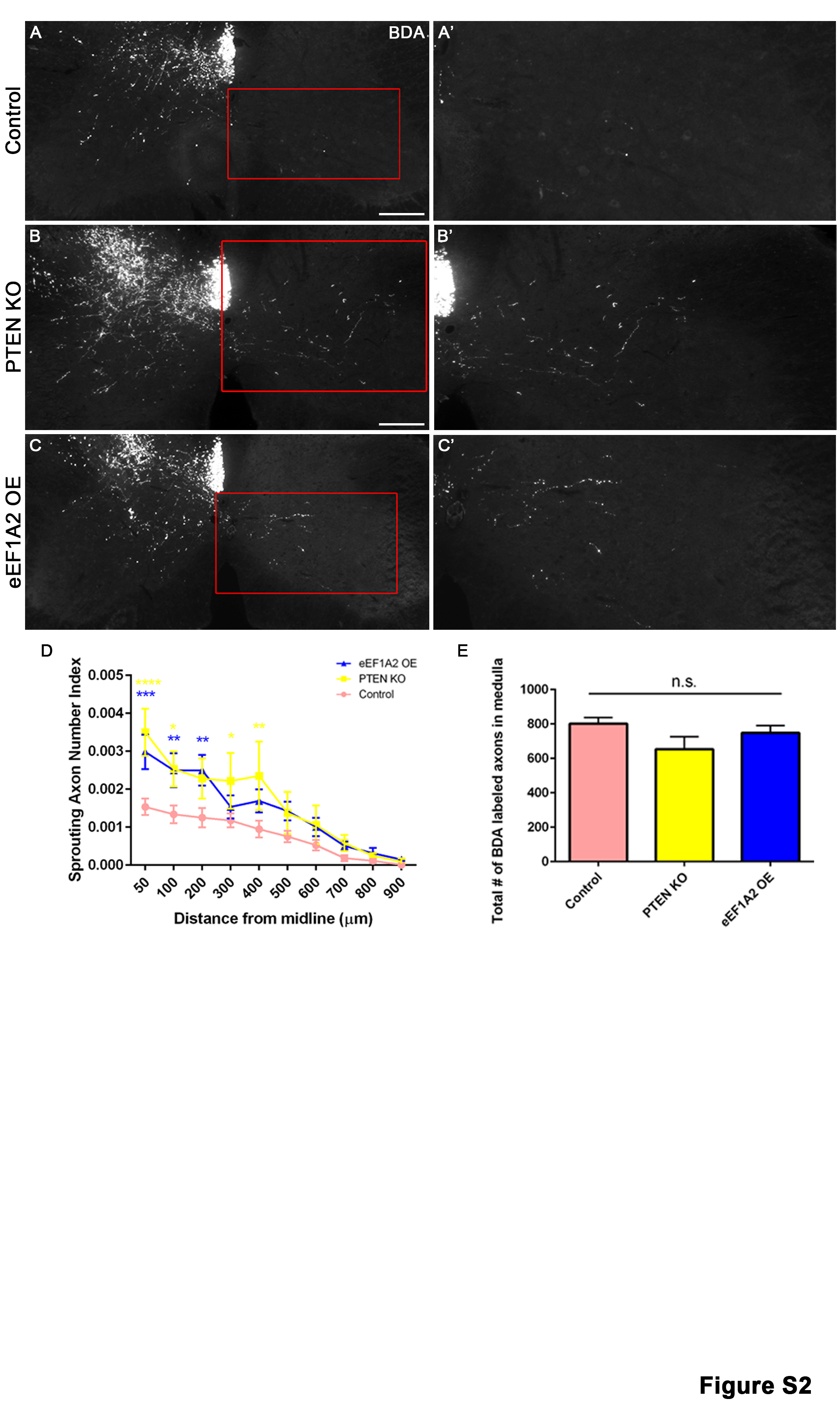
